# AmygdalaGo-BOLT: an open and reliable AI tool to trace boundaries of human amygdala

**DOI:** 10.1101/2024.08.11.607487

**Authors:** Quan Zhou, Bo Dong, Peng Gao, Jintao Wei, Jiale Xiao, Wei Wang, Peipeng Liang, Danhua Lin, the Chinese Color Nest Consortium, Xi-Nian Zuo, Hongjian He

## Abstract

Each year, thousands of brain MRI scans are collected to study structural development in children and adolescents. However, the amygdala, a particularly small and complex structure, remains difficult to segment reliably, especially in developing populations where its volume is even smaller. To address this challenge, we developed AmygdalaGo-BOLT, a boundary-aware deep learning model tailored for human amygdala segmentation. It was trained and validated using 854 manually labeled scans from pediatric datasets, with independent samples used to ensure performance generalizability. The model integrates multiscale image features, spatial priors, and self-attention mechanisms within a compact encoder-decoder architecture to enhance boundary detection. Validation across multiple imaging centers and age groups shows that AmygdalaGo-BOLT closely matches expert manual labels, improves processing efficiency, and outperforms existing tools in accuracy. This enables robust and scalable analysis of amygdala morphology in developmental neuroimaging studies where manual tracing is impractical. To support open and reproducible science, we publicly release both the labeled datasets and the full source code.

**Teaser:** An AI model enables accurate amygdala segmentation across ages and centers, supporting large-scale brain imaging studies.

## INTRODUCTION

Accurately segmenting the human amygdala, a small, structurally complex region, remains a major challenge in large-scale neuroimaging studies of brain development. The amygdala plays a central role in mood regulation, cognitive learning, and memory consolidation, and has been strongly implicated in behavioral pathologies - particularly during critical stages of socio-emotional maturation such as childhood and adolescence (*1–3*). Magnetic resonance imaging (MRI) has enabled detailed investigation of amygdala structure, facilitating the construction of developmental trajectories and the investigation of neurodevelopmental processes underlying various neuropsychiatric conditions (*4,5*). However, reliable volumetric analysis hinges on precise boundary-based segmentation—a task that remains technically challenging.

Conventional approaches have primarily relied on manual delineation of in vivo neuroimaging data, guided by histological atlases. While manual segmentation is considered the gold standard, it is time-consuming and infeasible for large-scale datasets. To address the limitations, researchers have developed automated segmentation tools based on probabilistic estimates (e.g., Markov random fields), Bayesian inference, and multi-atlas label fusion (*6–12*). These methods underpin popular tools like *FreeSurfer* and *volBrain*, which offer reasonable performance in whole-brain segmentation. However, these tools were not developed exclusively for amygdala segmentation. As a result, they struggle with boundary precision in this region, particularly in pediatric samples: *FreeSurfer* overestimates amygdala volume by 13–28%, while *volBrain* underestimates it by 36– 38% (*13*). These discrepancies underscore fundamental limitations of existing segmentation frameworks. The amygdala’s small size, low contrast with surrounding tissue, and complex anatomical context make it especially challenging to delineate accurately (*6, 11, 14*). Rule-based or atlas-driven models - such as those using probabilistic inference or multi-atlas fusion - struggle to accommodate such local variability, particularly in children and adolescents, where the amygdala is both smaller and still undergoing dynamic maturation. Consequently, systematic biases in volume estimation can compromise the precision and reliability of measuring amygdala, potentially undermining the validity of findings in developmental neuroimaging studies.

To address these challenges, we developed AmygdalaGo-BOLT, a boundary-aware deep learning algorithm trained on a large dataset of manually segmented pediatric amygdalae (*13*). The algorithm is built on a Transformer model (*15*) that emphasizes accuracy in boundary delineation. Rather than relying on predefined anatomical priors, it learns boundary features directly from data, enabling flexible adaptation to anatomical variability across development. In this study, we benchmarked AmygdalaGo-BOLT against widely used segmentation tools across internal and external neuroimaging cohorts, demonstrating its accuracy, stability, and generalizability for amygdala segmentation in developmental neuroimaging.

## RESULTS

### Performance on CKG-CCNC (Training Site)

Table 1 summarizes the comparative performance of BOLT3D against conventional segmentation algorithms within the CKG-CCNC test cohort. BOLT3D outperformed all other methods across multiple key metrics, demonstrating superior segmentation accuracy and boundary precision. It achieved a Dice coefficient of 0.904, notably higher than U-Net3D (0.851), U-Net3D++ (0.787), and ResU-Net3D (0.841), indicating greater overlap with manual annotations. A similar pattern emerged for the Jaccard Similarity (JS) coefficient, with BOLT3D reaching 0.825. In terms of boundary accuracy, BOLT3D achieved the lowest 95th-percentile Hausdorff Distance (HD95) of 1.097, indicating superior alignment with anatomical ground truth compared to U-Net3D (1.565), U-Net3D++ (1.963), and ResU-Net3D (1.669). The average symmetric surface distance (ASSD) further demonstrated that BOLT3D’s segmentation output aligned more closely with reference annotations compared to other methods. Evaluation of detection performance further highlighted BOLT3D’s advantage. It achieved a precision of 0.883 and recall of 0.929, resulting in a high F1-score of 0.905, reflecting its ability to balance false positive suppression with high true positive detection. Finally, BOLT3D attained a relative volume difference (RVD) of just 0.056, over 40% lower than the next-best method, underscoring its accuracy in volumetric estimation.

**Table 1.** Performance evaluation on different methods of tracing amygdala in the CKG-CCNC cohort.

|  |  |  |  |  |  |  |  |  |
| --- | --- | --- | --- | --- | --- | --- | --- | --- |
| Method | Dice | JS | HD95 | ASD | ASSD | Precision | Recall | RVD |
| FreeSurfer | 0.776 | 0.636 | 2.530 | 1.093 | 0.954 | 0.732 | 0.832 | 0.142 |
| U-Net3D | 0.851 | 0.743 | 1.565 | 0.746 | 0.693 | 0.774 | 0.951 | 0.239 |
| U-Net3D++ | 0.787 | 0.651 | 1.963 | 1.064 | 0.982 | 0.658 | 0.985 | 0.512 |
| ResU-Net3D | 0.841 | 0.728 | 1.669 | 0.806 | 0.736 | 0.754 | 0.957 | 0.279 |
| BOLT3D | **0.904** | **0.825** | **1.097** | **0.483** | **0.467** | **0.883** | **0.929** | **0.056** |

Fig. 1 presents segmentation results from multiple algorithms on a representative participant, demonstrating that BOLT3D most closely aligns with manual reference standard in both volumetric estimation and boundary delineation. Notably, the amygdala delineated by FreeSurfer deviates markedly from both the morphological and manual segmentation. The presence of multiple empty regions indicates an inability to accurately capture the structural integrity of the amygdala. A closer inspection of the periphery suggests that FreeSurfer exhibits numerous irregularities when compared with the manual reference standard. Although Unet produces segmentations that are morphologically more consistent with manual references, volumetric evaluation reveals substantial overestimation. In contrast, BOLT3D produces segmentation results that more closely match manual delineations in both overall shape and boundary accuracy. Notably, the superposition and edge correspondence in BOLT3D outputs demonstrate minimal deviation from reference, indicating precise volume estimation and structural fidelity.

**Fig. 1.**
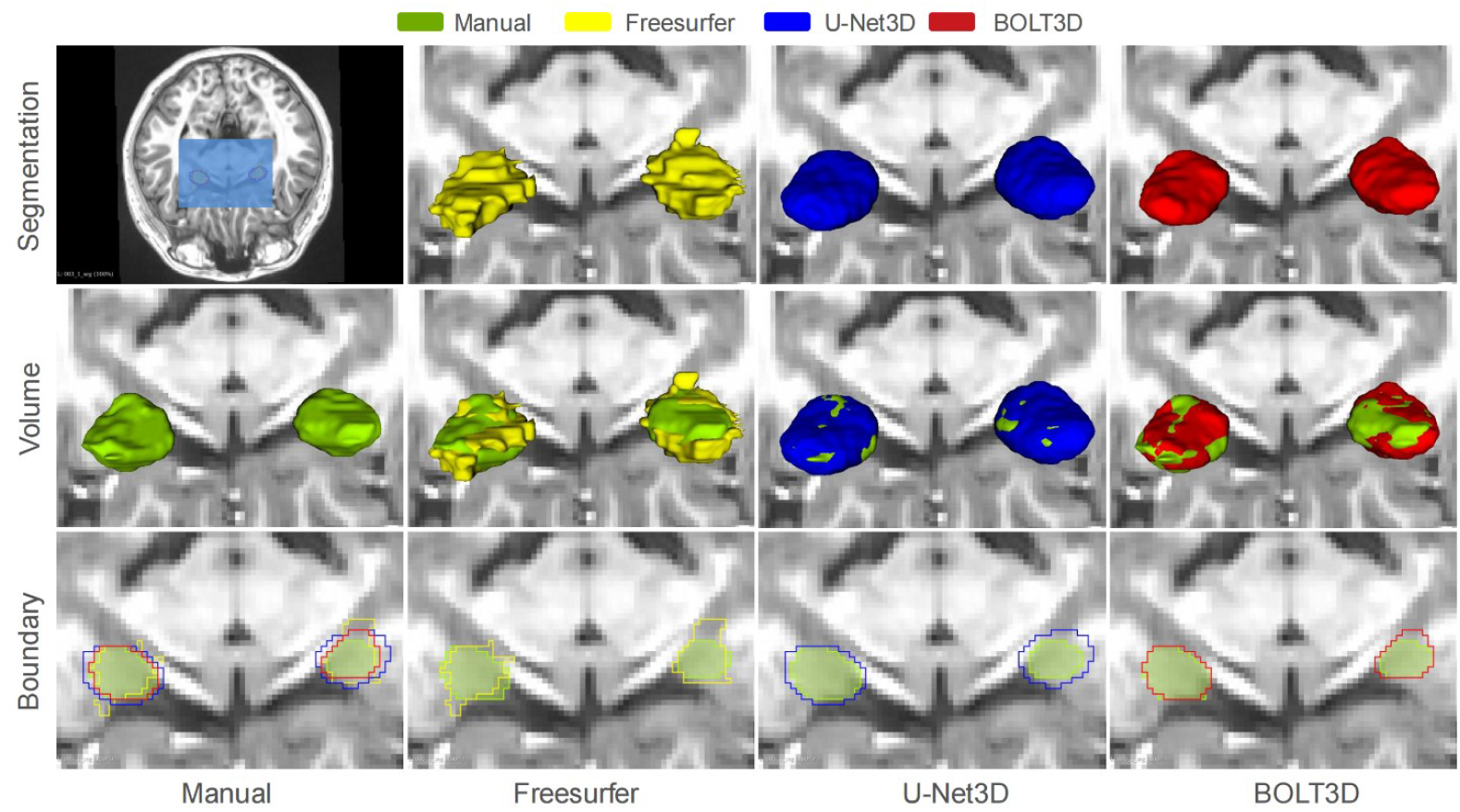
Comparative visualization of amygdala segmentation in a representative participant. The segmentation results of different methods illustrate the preservation of structural integrity (middle row) and the precision of boundary delineation (bottom row). Manual segmentation (left column) serves as the reference standard. The proposed BOLT3D method (right column) shows superior performance in both volumetry and boundary definition, outperforming FreeSurfer (second column) and U-Net3D (third column).

The generalizability of BOLT3D was first validated on an dataset from BNU-CAC, an independent imaging center not included in model training, comprising 116 children. As shown in Table S1, BOLT3D maintained strong performance despite distributional shifts, achieving a Dice coefficient of 0.845 and a Jaccard similarity (JS) of 0.733. Distance-based evaluations further confirmed accuracy, with HD95 and ASD values of 1.594 and 0.654, respectively. The average symmetric surface distance (ASSD) of 0.666 exhibited less variability than that observed with U-Net, suggesting more stable boundary delineation. In terms of detection performance, BOLT3D attained a precision of 0.933 and a recall of 0.775, effectively balancing false-positive suppression and true-positive identification. The relative volume difference (RVD) was 0.168, indicating low deviation in volumetric estimation. These indicators suggest the performance of BOLT3D was not compromised significantly, despite the distribution variations encountered at new centers. In addition, this technique significantly outperformed comparable algorithms, thereby affirming the multi-center accuracy and consistency offered by integrating the consistency module. Here, we presented segmentation results from 116 unseen participants in BNU-CAC, compared to the 347 observed scans from CKG-CCNC (see Fig. 2). Since BNU-CAC participants originate from different data distribution and were not included in model training, the overall performance on this dataset is lower compared to the CKG-CCNC scans. This performance gap likely arises due to variations in feature distributions, inter-data noise, or other unaccounted factors that affect segmentation accuracy. Despite this decline in overall performance with unseen data, BOLT3D significantly outperforms the baseline model, demonstrating strong generalization capabilities.

**Fig. 2.**
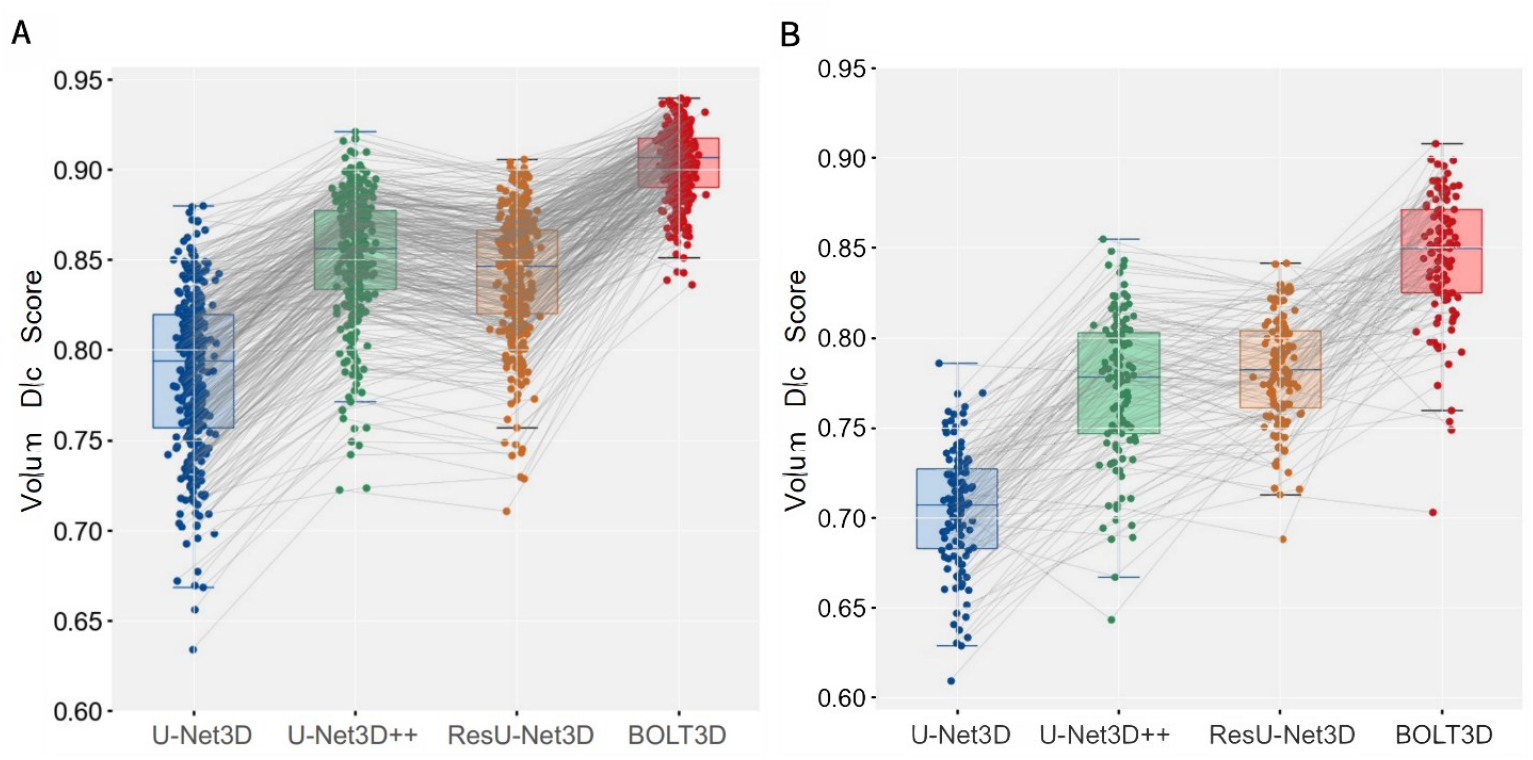
Volume dice analysis of the overall amygdala structure for participants from the CKG-CCNC (A) and BNU-CAC (B) cohorts. Since the CKG-CCNC data were used during training, the overall performance was slightly higher than for the unseen BNU-CAC dataset for all deep-learning network methods. However, the proposed BOLT3D model achieved a dice score above 0.8 for both cases.

To further evaluate cross-site robustness, the model was tested on the ZJU-MTC dataset, which includes three “traveler” participants scanned at ten different imaging centers (Fig. 3). BOLT3D consistently outperformed U-Net3D across all sites, with Dice improvements ranging from 0.02 to 0.07. These results highlight BOLT3D’s resilience to multi-center heterogeneity and support its applicability to large-scale, multi-center neuroimaging studies.

**Fig. 3.**
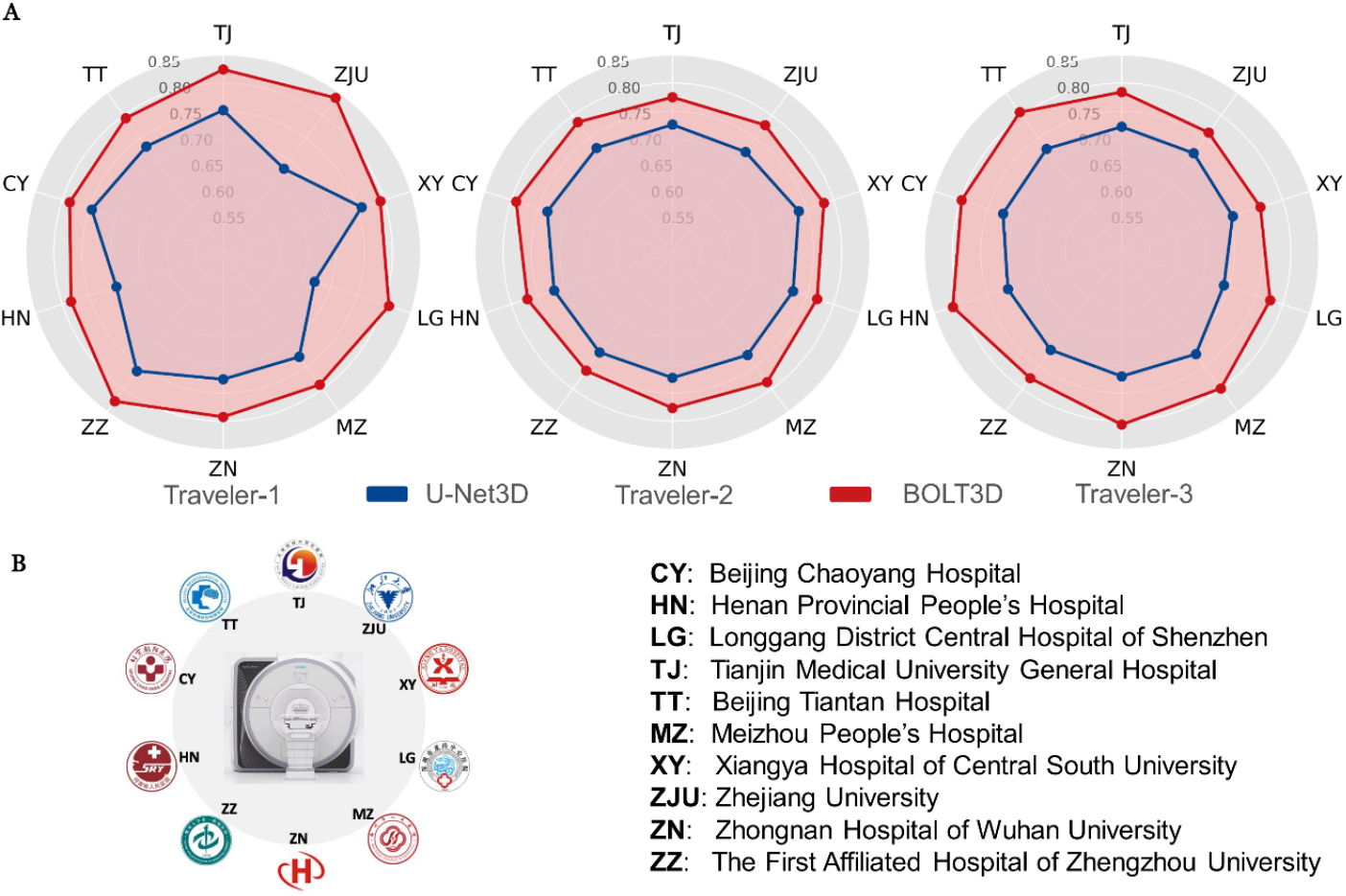
A comparison of amygdala segmentation outcomes for itinerant volunteers from ZJU-MTC. (A) The proposed BOLT3D methodology (depicted in red) consistently demonstrated superior performance relative to U-Net3D (depicted in blue) across all centers for each participant, thereby underscoring its robust cross-center generalization efficacy. (B) All centers were equipped with identical model 3T scanners, geographically dispersed across China, with their respective names and abbreviations provided on the right.

### Ablation Study in ZJU-MTC

We conducted ablation experiments on the ZJU-MTC dataset to evaluate the contribution of key BOLT3D components to segmentation accuracy and generalization. As shown in Fig. 4A–B, the model variant incorporating the MGAC backbone (but excluding the SAM module) yielded substantially higher Dice scores and lower ASSD values compared to U-Net3D and FreeSurfer, across both volumetric and boundary metrics. These results reflect MGAC’s ability, via a Transformer-based multi-granularity attention framework, to capture both global context and local anatomical detail, thereby improving segmentation accuracy and structural fidelity.

**Fig. 4.**
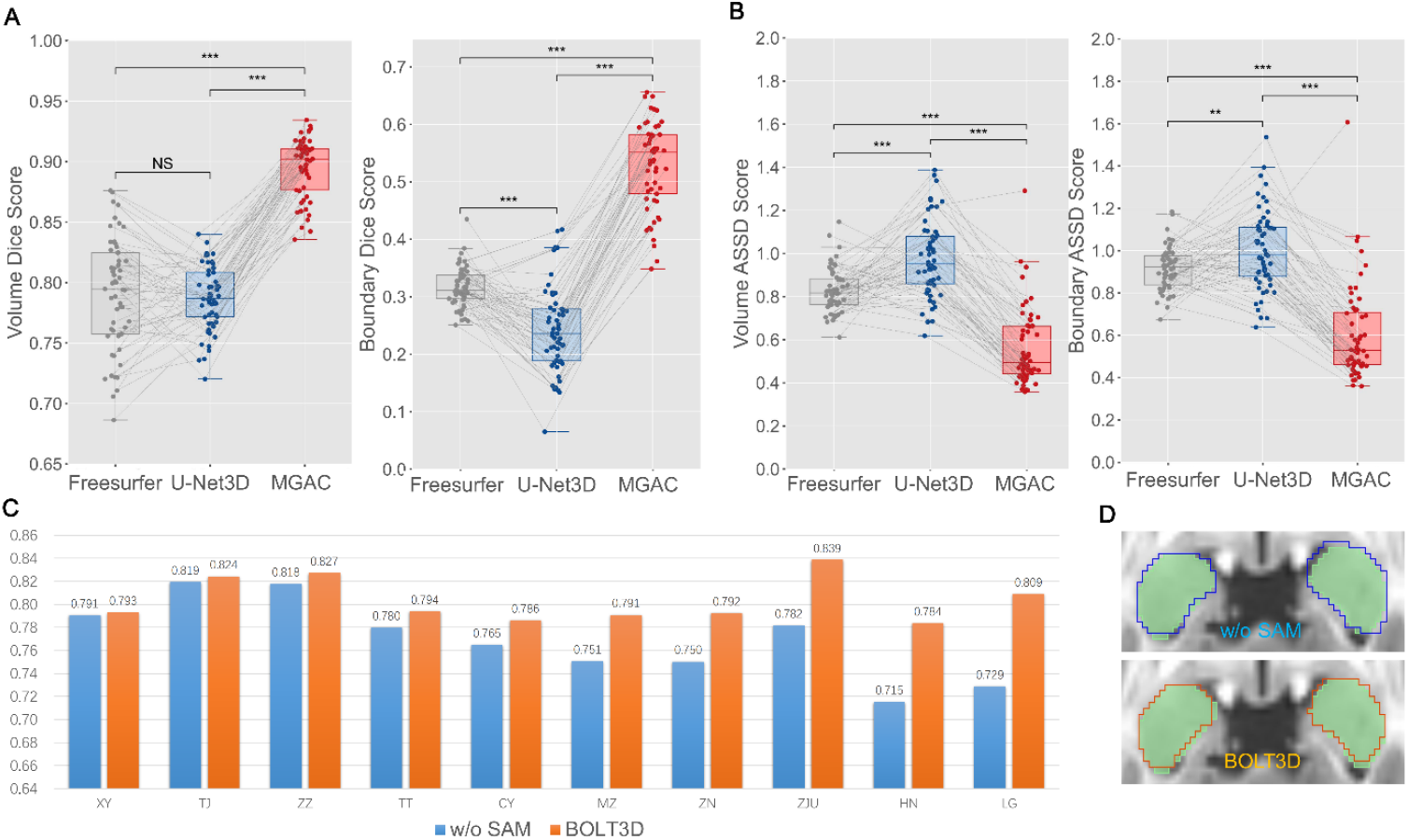
Ablation study evaluating the contributions of MGAC and SAM modules. (A–B) The MGAC module improves Dice and ASSD scores compared to U-Net3D and FreeSurfer. (C) Integration of the SAM module enhances robustness across imaging centers in the ZJU-MTC dataset. (D) Visual examples illustrate improved boundary fidelity with SAM.

We then assessed the SAM module’s effect on cross-site consistency by comparing segmentation performance across ten imaging centers. Integration of SAM into the full BOLT3D pipeline led to consistent performance under diverse scanning conditions gains over the MGAC-only variant (Fig. 4C), demonstrating its utility in mitigating inter-site variability. Qualitative comparisons further illustrated improved boundary adherence following SAM-based refinement (Fig. 4D).

### Foster Fast and Valid Large-scale Amygdala-centered Neuroscience Research

To assess segmentation validity, we evaluated the Pearson correlation between automated and manual amygdala volume estimates using 347 internal validation scans from the CKG-CCNC dataset (excluding the training set). As shown in Fig. 5, BOLT3D achieved the highest correlation with manual segmentation for both the left (R = 0.74) and right (R = 0.70) amygdalae, outperforming U-Net3D, U-Net3D++, ResU-Net3D, and FreeSurfer (R = 0.46–0.61). This strong linear association demonstrates BOLT3D’s superior accuracy in volumetric quantification. In addition, the narrow scatter distribution observed with BOLT3D highlights its within-subject consistency and stability across individuals. To further quantify agreement and systematic deviation, Bland–Altman analysis was conducted on the same dataset (Fig. 6). BOLT3D showed the narrowest limits of agreement (–0.66 to +0.35 mm^3^) and the smallest mean difference from manual labels (–0.152 mm^3^), indicating minimal bias and high consistency. In contrast, alternative methods exhibited substantially greater overestimations, with average differences ranging from – 0.647 to –1.874 mm^3^. These discrepancies were up to 12.329 times larger than those observed with BOLT3D, emphasizing its advantage in volumetric measurement precision.

**Fig. 5.**
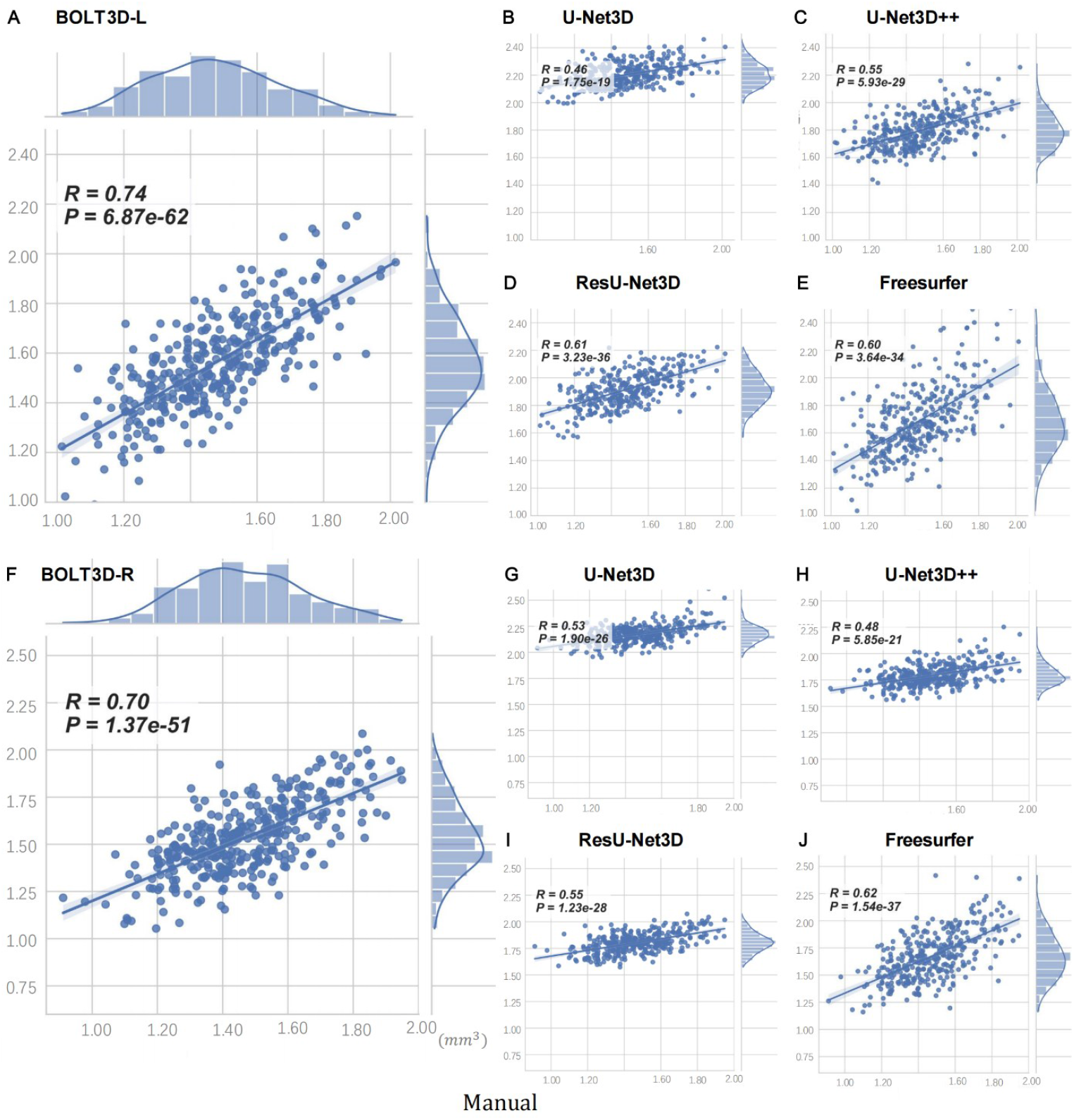
A correlation analysis of amygdala volume as determined by segmentation algorithms compared with manual reference standards. The dataset included 347 internal validation scans, with the results of manual segmentation depicted on the horizontal axis. Among the evaluated methods, the proposed BOLT3D algorithm achieved the highest correlation with manual references for the left (A) and right (F) regions of the amygdala, signifying higher consistency in volumetric measurements. In contrast, correlation coefficients for alternative segmentation tools, including U-Net3D (B, G), ResU-Net3D (C, H), U-Net3D++ (D, I), and FreeSurfer (E, J), fell within a range of 0.46 to 0.60. The volume distributions derived from each algorithm are represented on the respective vertical axes.

**Fig. 6.**
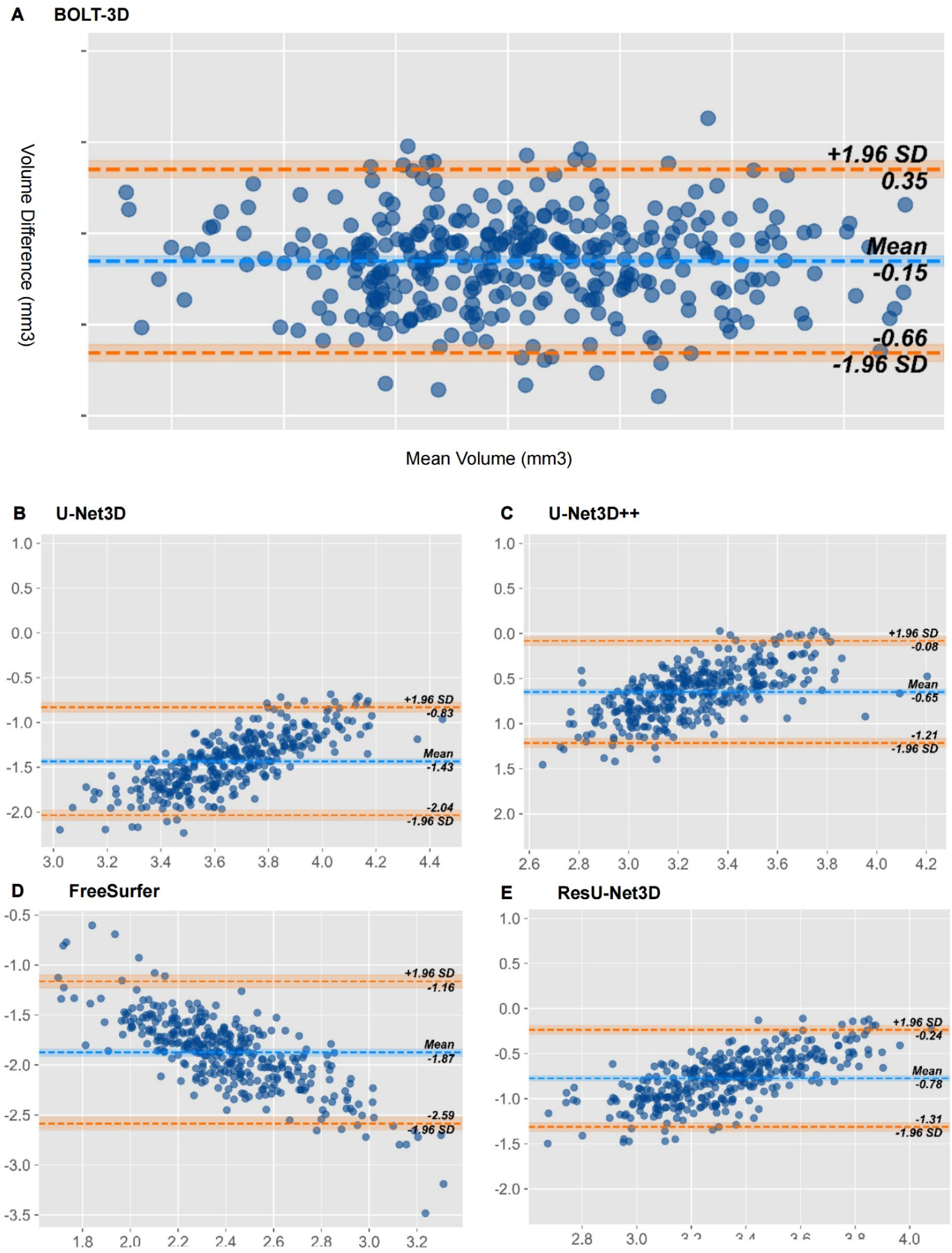
Bland–Altman analysis of volumetric agreement between automated methods and manual reference. Bland–Altman plots were used to assess the agreement between volume estimates from different automated segmentation algorithms and the manual reference standard across 347 left amygdala scans. The central dashed blue lines indicate the mean differences, and the dashed yellow lines represent the limits of agreement (±1.96 standard deviations from the mean). Each blue dot corresponds to an individual subject.

To evaluate the capability of each algorithm to capture developmental trajectories, we plotted amygdala volume growth curves across age using algorithm-derived segmentations (Fig. 7). These were benchmarked against manually annotated volumes, considered the reference standard (*13*). Both manual and automated segmentations revealed parallel growth patterns for boys and girls, with boys consistently exhibiting larger amygdala volumes throughout development. Manual tracing indicated a linear increase in bilateral amygdala volume from childhood to early adulthood, a trend closely mirrored by BOLT3D, which employed multi-feature fusion. This was supported by the estimated degrees of freedom (edf) reported in Table S2. Although the bilateral amygdala volumes predicted by BOLT3D were slightly larger than those of manual annotations, the scatter plot exhibits a more confined aggregation among age cohorts. In contrast, FreeSurfer produced an inverted U-shaped trajectory that deviated markedly from the linear pattern observed in both manual and BOLT3D results. Thus, the scatter plots reveal an increase in variability between age cohorts, with bilateral amygdala volumes exceeding those derived from both manual annotations and BOLT3D. This outcome demonstrates significant advantages for the fitting of growth curves compared to other algorithms, attesting to the robustness and accuracy of BOLT3D in considering gender- and age-related changes when estimating amygdala volume. Compared to conventional and other advanced algorithms, this technique excels at capturing subtle changes in growth curves, not only in terms of overall adaptation efficiency but also in terms of accurately predicting curve details and particular features. These findings improved our understanding of amygdala development over time with respect to gender differences and age-related factors.

**Fig. 7.**
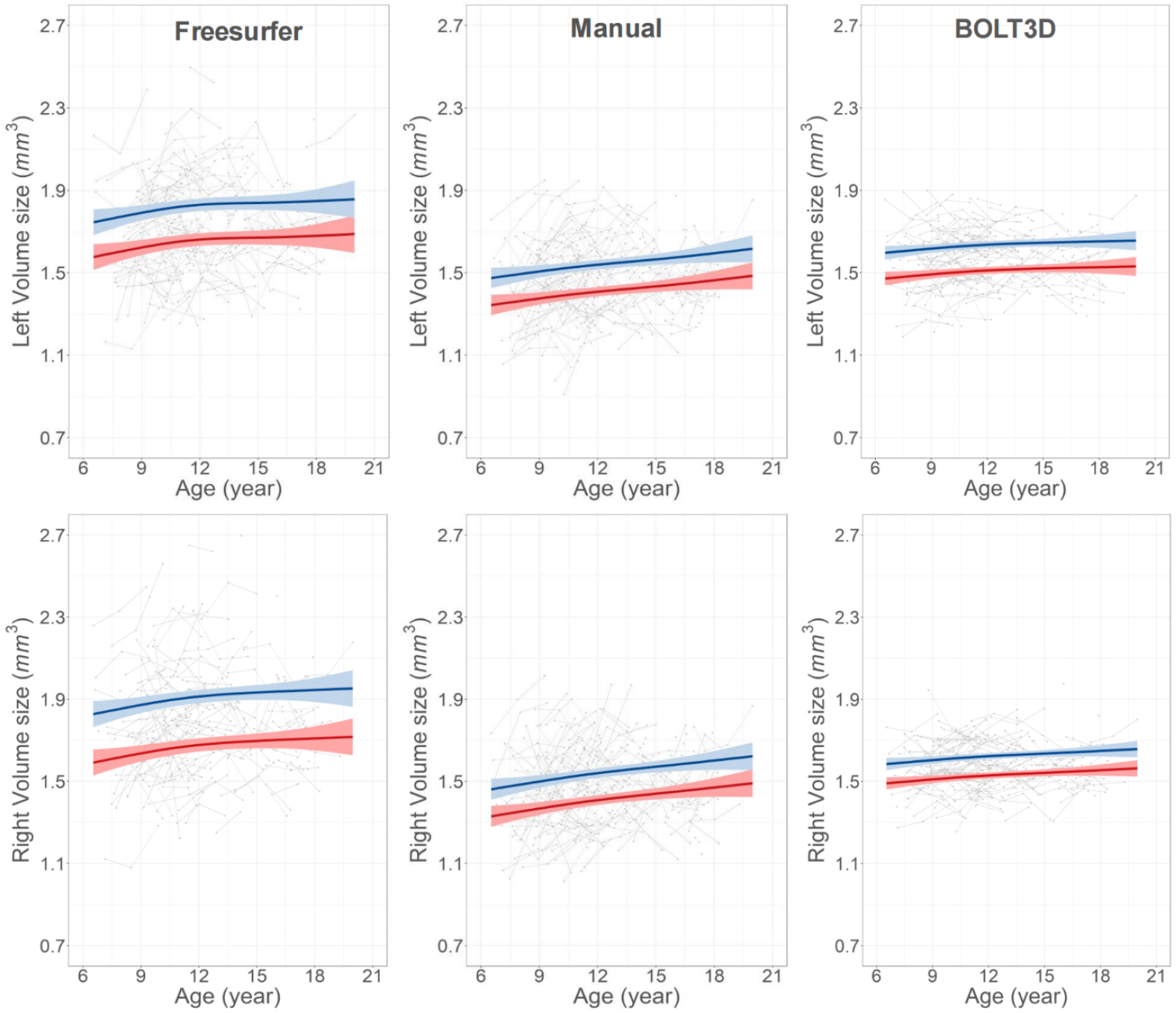
The developmental trajectory of the amygdala, as delineated by our model and comparative methodologies. The FreeSurfer results (left) exhibit an inverted U-shaped curve, indicative of a non-linear growth pattern. Conversely, the growth curves predicted by our BOLT3D model (right) exhibit a striking agreement with the linear trajectory observed in the manual tracing method (middle). Blue denotes trajectories for males, while red represents trajectories for females. These trajectories are enveloped by shaded 95% confidence intervals. Individual scatter points represent segmentation outcomes from distinct cases, while the lines connecting these points indicate results acquired from multiple scans of the same participant.

### AmygdalaGo-BOLT3D Model Availability

The AI training platform supporting this work was provided by High-Flyer AI (Hangzhou High-Flyer AI Fundamental Research Co., Ltd.). We deploy the optimal model parameters to be applied to any user-specified dataset in the interest of open science as well as provide tutorials and source codes at the Science Data Bank (https://ccnp.scidb.cn/en).

## DISCUSSION

Deep-learning methods of tracing boundaries of amygdala are highly valuable for reproducibility by providing standardized, automated pipelines that minimize the variability introduced by manual tracing or traditional atlas-based methods. In open neuroscience, where large-scale datasets are shared across institutions, AI-driven segmentation ensures consistent and reproducible annotations, making results more comparable across studies. In the present work, we develop an open-source AI tool, AmygdalaGoBOLT3D, which allows researchers to collaborate, validate, and refine models using publicly available datasets, fostering transparency and innovation. Traditional segmentation techniques, whether manual or atlas-based, are time-consuming and prone to inter-rater variability, especially for small structures like the amygdala. A deep-learning model such as AmygdalaGO-BOLT3D trained on large datasets can provide 1) higher segmentation accuracy, especially for challenging cases with low-contrast boundaries; 2) faster processing, reducing the time required for segmentation from hours to seconds; 3) improved generalization, allowing them to adapt to different populations, imaging protocols, and scanner types. Another example of AI-based segmentation using multi-atlas priors and 3D convolutional networks (*32*) has demonstrated superior performance over traditional methods.

Our AmygdalaGo addressed the intricacies posed by low contrast and structural complexities that distinguish the amygdala from adjacent tissues. A boundary contrastive learning algorithm was efficient to align model-predicted boundaries more accurately with a ground truth, culminating in more precise segmentation across a wide range of subject ages, ranging from childhood and adolescence to adulthood. It also maintained remarkably consistent performance when deployed in multi-center adult datasets, underscoring exceptional generalization proficiency and robust performance across varied data sources and contexts. This transformer could be a valuable new tool in the field of cognitive neuroscience. For example, previous studies have identified a relationship between the size of social networks and the volume of the amygdala (*33–35*).

Recent advances in open neuroimaging data have significantly increased our understanding of the intricate structures and networks within the human brain (*36*). The aggregation of images from various large-scale data repositories has significantly improved statistical power, allowing for the discernment of subtle variability within the human amygdala (*37,38*). However, there is a pressing need for new research methodologies to enhance the computational efficiency and practical applicability of these automated extraction tools (*13,39*). Global imaging collaborations have been instrumental in the evolution of population neuroscience (*40,41*), driven by extensive MRI initiatives (*42*), which have been crucial for driving innovations in analytical techniques and translational research (*43*). As an open-source deep-learning segmentation tool, AmygdalaGO-BOLT3D democratizes access to high-quality neuroimaging analysis, making it possible for researchers without extensive expertise in neuroanatomy to conduct high-precision studies. This is crucial for 1) low-resource institutions that may lack access to expert manual tracers; 2) large-scale, multi-site collaborations where standardized, automated segmentation ensures data harmonization; 3) early-career researchers and students who can leverage open-source AI tools without requiring specialized training in segmentation techniques. By integrating deep learning into open neuroscience platforms, the AmygdalaGO-BOLT3D pipeline can be widely distributed, continually improved, and used globally, accelerating discoveries in neurodevelopment, psychiatric disorders, and neurodegenerative diseases.

We note that the present study lacks a detailed subdivision of the amygdala, specifically finer-grained nuclei labeling. Future work will focus on providing granular annotations and establishing a computational baseline to support a more nuanced analysis of the amygdala. Unlike static atlas-based segmentation tools, AmygdalaGO-BOLT3D improves over time by leveraging crowdsourced data and federated learning approaches. This means that as new datasets become available, its model can be updated, enhancing its robustness and adaptability. Community-driven platforms such as OpenNeuro (https://openneuro.org/), NeuroVault (https://neurovault.org/), and ScienceDB (https://ccnp.scidb.cn/en) can serve as repositories for the model training and validation, ensuring that this AI-driven technique remains cutting-edge and inclusive.

AmygdalaGo-BOLT, a deep-learning method for boundary tracing in human brain images, offers a scalable, accurate, and open-access solution for amygdala segmentation. It aligns with the principles of open neuroscience in four keyways: 1) Reproducibility - AI-driven segmentation ensures consistent and replicable results across studies; 2) Efficiency - Faster and more precise segmentation compared to manual tracing; 3) Accessibility - Enables global participation in neuroimaging research; 4) Adaptability - The model improves over time with diverse datasets. Unlike conventional approaches, BOLT3D incorporates an MGAC module, specifically designed to integrate global and local semantic features. Additionally, its boundary contrast learning strategy refines edge delineation, addressing the persistent challenge of distinguishing the amygdala from adjacent structures. These innovations establish AI-driven segmentation as a transformative tool for large-scale brain imaging, advancing more inclusive, transparent, and impactful discoveries in amygdala neuroscience.

## MATERIALS AND METHODS

This section provides an overview of the datasets, model architecture, training procedures, and evaluation protocols used in the development and validation of AmygdalaGo-BOLT3D. We first describe the MRI datasets employed for model training and validation. Next, we detail the network architecture, including its multi-granularity adaptive collaboration, boundary contrast learning, and transformer-based refinement modules. Subsequently, we explain the training configuration and implementation settings. Finally, we present the evaluation metrics utilized to assess segmentation performance.

### Datasets

#### Training Dataset

The AmygdalaGo-BOLT3D model was trained using data from the Chongqing cohort of the Chinese Color Nest Cohort (CKG-CCNC), part of the CCNP development project (*16–18*). This cohort included 198 baseline participants aged 6–17 years, from whom a total of 427 T1-weighted MRI scans were acquired. To ensure balanced age distribution and demographic representativeness, 140 participants were selected from the full pool. For each, one high-quality scan was retained after rigorous visual inspection to exclude motion artifacts and anatomical abnormalities. From these, 80 scans were randomly selected for model training (Table 2). Manual boundary tracing of the amygdala was performed according to standardized protocols (*13*).

**Table 2:**
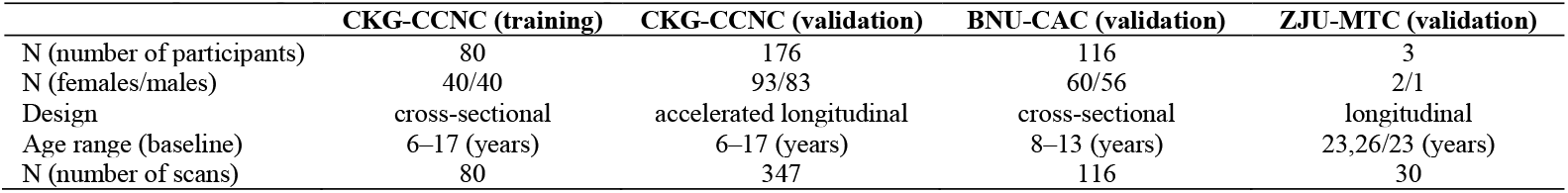
Sample demographic information and sample characteristics of the three testbed datasets.

#### Validation Datasets

Model validation was conducted using three datasets. First, the remaining 347 MRI scans from the full CKG-CCNC sample (excluding the 80 used for training) were used for internal validation. Second, external validation was performed using brain MRI data from two independent cohorts: the Beijing Normal University Childhood Adversity Cohort (BNU-CAC), comprising 116 children aged 8–13 years scanned once (*19*), and the Zhejiang University Multi-center Traveler Cohort (ZJU-MTC), which included three adult participants scanned across ten different MRI centers using harmonized protocols (*20*). In all validation datasets, the amygdalae were manually segmented using the same boundary tracing procedures as in the training set to ensure consistency. During testing, input image dimensions were down-sampled to 48 × 64 × 80 to minimize computational demands. No post-processing optimization techniques were employed, thereby ensuring an objective assessment of model performance.

### Model Architecture

#### Overall Framework

We implemented AmygdalaGo-BOLT in three stages: 1) robust global localization, 2) precise delineation, and 3) generalizable segmentation. When examining the structure of the amygdala, the observer must first capture the entire cerebral region and subsequently estimate the approximate position of the amygdala within the image. This estimate relies on prior knowledge of limbic system anatomy and considers adjacent structures such as the hippocampus. After this initial phase, the observer leverages a detailed understanding of internal amygdala morphology, gray matter intensity, and other pertinent information used to execute a full regional delineation. The segmentation process is then enhanced using a transformer-based algorithm employed to discern boundary contrast. It aims to improve both the robustness of global amygdala localization and the precision of boundary delineation, thereby improving inter-individual consistency. This process includes the following key elements:

1. Comprehensive detection of amygdala positioning in T1-weighted (T1w) MR images was achieved by employing a transformer-based network. This model (BOLT3D) constructs a multi-granularity adaptive collaboration (MGAC) architecture, which integrates a global coarse-grained perceptual module. It determines the global semantics of the amygdala structure, thereby enhancing positional sensitivity. Since intricate details are often neglected in global representations, a local fine-grained enhancement module was developed to increase the perception and retention of detailed information. Coarse-grained global and fine-grained local data were further harmonized by introducing a multi-granular adaptive fusion module. This allowed BOLT3D to concurrently consider differences between the entire amygdala and adjacent structures, as well as similarities within the amygdala itself. It facilitates the recognition of global positioning characteristics and local disparities, thereby improving the accuracy and robustness of amygdala structure location estimates. The overarching design objective is then to perform a more precise and exhaustive detection of amygdala tracing contours.
2. A novel strategy is introduced for learning boundary contrast, to address dependency challenges for manual segmentation boundaries and enhance the congruence between predicted and actual delineations. It improves the delineation of limbic structures from adjacent tissues by amplifying characteristic differences between the amygdala and surrounding tissues, while minimizing intra-amygdala variations. Consequently, this step refines the alignment between automated boundary segmentation and manual segmentation outcomes, enabling a more precise demarcation of amygdala dimensions.
3. A modified Segment Anything Model (SAM) is introduced to refine coarse predictions and enhance the generalizability of BOLT3D. Coarse outputs from the MAGC module are sampled as point cues and fed into the 3D Prompt Encoder, which integrates anatomical priors to enhance segmentation accuracy. To adapt SAM for amygdala segmentation, three key modifications are implemented: (1) a lightweight Volumetric Feature Encoder (VFE) to extract MRI-specific volumetric features, (2) a 3D Prompt Encoder (PE) to refine spatial embeddings and anatomical representations, and (3) a Volumetric Mask Decoder (VMD) to enhance segmentation precision by processing the refined feature representations. These adaptations enable SAM to effectively manage the amygdala’s small size, improve boundary delineation via contrastive learning, and enhance generalization across diverse subjects. By integrating these domain-specific refinements, the model achieves greater segmentation accuracy and robustness.

#### Multi-Granularity Adaptive Collaboration

Global and local semantic features were initially derived from the original MR image data using a multi-granularity semantic generation module, as depicted in Fig. 8B. Specifically, for T1w images *I*, the module exploited local induction capabilities by employing two 3D convolutions to extract primary local features *F*_*l*_. This approach facilitated the computation of relationships among feature points, thereby enabling the extraction of global semantic information from regions exhibiting high responses to initial position features. Subsequently, preliminary global features *F*_*g*_ were constructed as follows. A feature map *F′* of dimensions *d* × *H* × *W* was derived from the input image *I* using a single convolutional layer and subsequently projected onto a weighting map via two convolutional layers. The channel count in the weight map was given by *hw* (*h* ≤ *H, w* ≤ *W*), where *h* and *w* represent the dimensions of a predefined global semantic map. The weight map was then flattened for the Softmax computation, and the resulting weights were used to aggregate semantic information within the *F′* mapping through matrix multiplication, producing the final *F*_*g*_ result. In the following illustrations, dimensions are presented in an abbreviated form for brevity.

**Fig. 8.**
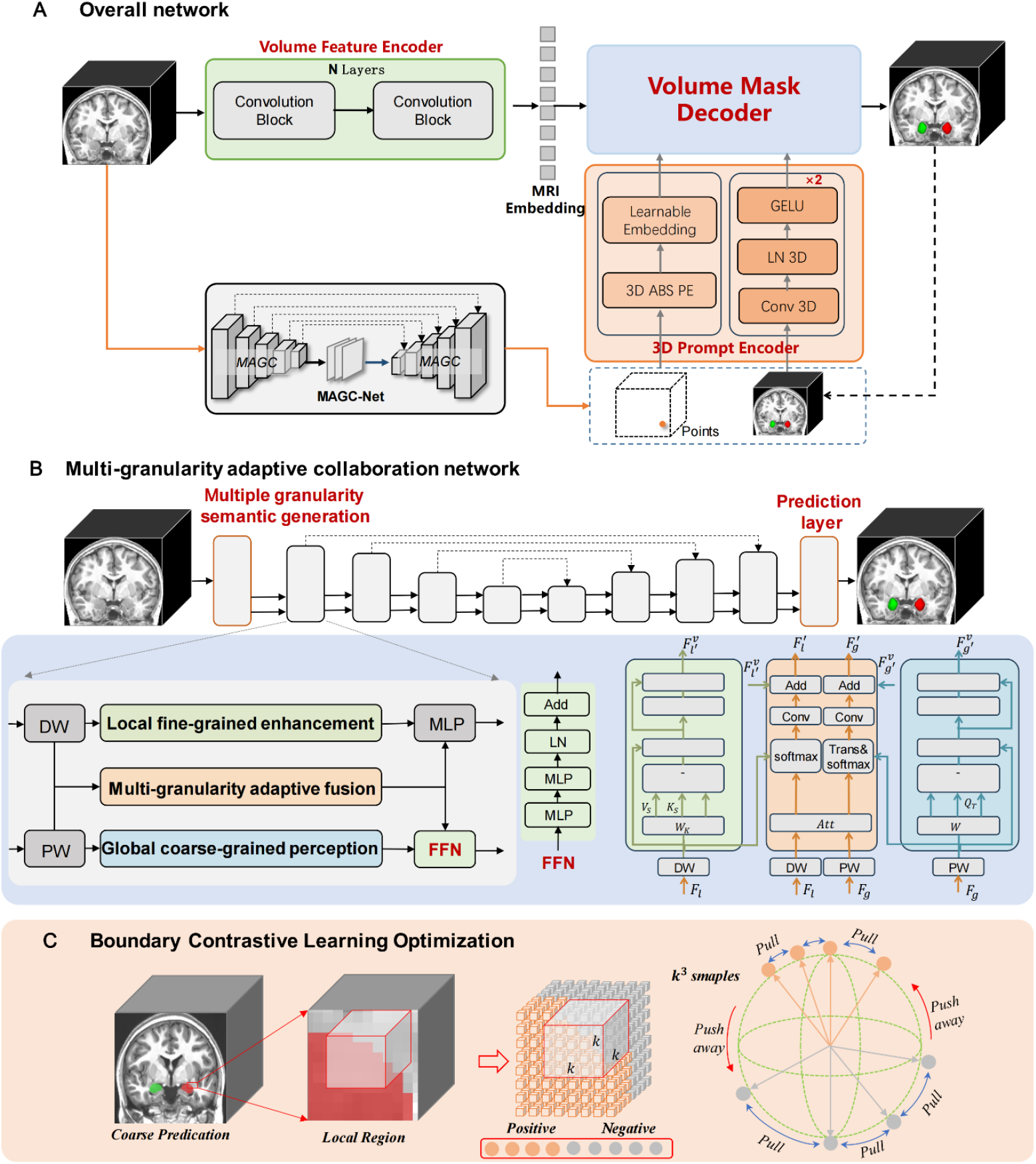
A schematic diagram of the AmygdalaGo-BOLT architecture. (A) The overall model structure consists of two principal components: a MAGC-Net (responsible for initial segmentation) and an SAM. The latter is tasked with the refinement of segmentation results, including three essential components: a lightweight Volume Feature Encoder (VFE), a 3D Prompt Encoder (PE), and a Volumetric Mask Decoder (VMD). The MAGC network produces preliminary segmentation results and strategically samples voxels, which are then fed into the SAM prompt encoder to guide the refinement process. (B) The MAGC block, a key network module designed to address challenges posed by the size of the amygdala, ensures the extraction of accurate semantic features. (C) The included edge contrast learning optimization strategy, incorporated to further augment edge delineation and enhance overall segmentation precision.

The MGAC module includes three components: a global module for coarse-grained perception, a local module for fine-grained enhancement, and an adaptive module for multi-granularity fusion, as depicted in Fig. 8C. These components independently optimize global and local features, which are interactively integrated to enhance model performance. Fig. 8B illustrates the positioning of the adaptive MGAC module within the overall framework, while Fig. 8C details the corresponding internal structure. Each MGAC module receives two inputs: global coarse-grained (*F*_*g*_) and local fine-grained (*F*_*l*_) semantic characteristics. Significant discrepancies in semantic distributions between these two sets of characteristics were mitigated by using BatchNorm3D to normalize the inputs. A linear position projection from previous work was then mapped based on the positional relationships of the elements, ensuring the arrangement of tokens within the transformer remained unchanged while preserving their positional correlations (*21*). However, this approach neglects local structural information, which is vital for image data. Although integrating positional encoding enables the transformer to learn positional relationships, such encoding from randomly initialized parameters requires extensive training data, which poses a challenge for medical image analysis. Traditional positional encoding data were then substituted with convolutional layers, thereby incorporating the inductive bias inherent in convolutions. This substitution facilitated the preservation of local structural information intrinsic in the images while avoiding the limitations of positional encoding.

A three-dimensional (3D) depth-wise convolution strategy was employed to convolve relationships among local neighborhoods within fine-grained semantic features, thereby introducing visual bias. In contrast, 3D pointwise convolutions were used to calculate global coarse-grained semantic features. The results, denoted as *F*_*g*_ and *F*_*l*_, were subsequently flattened into a one-dimensional (1D) sequence to facilitate Transformer processing. This transformation involved converting *F*_*g*_ into a global query vector 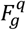 and a global content vector 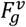. Similarly, *F*_*l*_, which encapsulates detailed information, was transformed into a local index vector 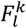 and a local content vector 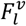 through analogous operations. In contrast, the internal architectures of the global coarse-grained perception module and the local fine-grained enhancement module remained consistent, as illustrated in Fig. 8B. These modules are primarily comprised of two components: a multi-head self-attention mechanism and a feedforward neural network. The *Z*_*l*_ term initially replaced the input in the multi-head self-attention mechanism and was then expanded into a 1D sequence. A fully connected layer served as the transformation layer, converting the input to a query vector, an index vector, and a content vector. The multi-head attention matrix could subsequently be determined as:

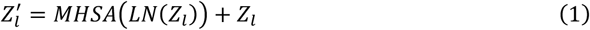

where, the feedforward layer further refined the relationships among voxel features, while integration with residual mechanisms augmented the data flow. This approach preserved original features and mitigated vanishing gradients, as described by:

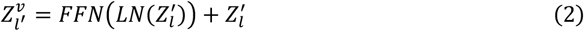

In this context, *MHSA*(·) denotes a multi-head self-attention mechanism, *LN*(·) signifies layer normalization, and *FFN*(·) represents a feedforward neural network. The global coarse-grained perception module and the local fine-grained enhancement module employed Eqs. 1 and 2 to transform the input global content vector 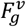 and the local content vector 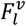, respectively. The resulting updated global content vector 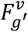 and the local content vector 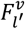 were obtained upon completion of the processing step. The correlation matrix *Att* was then derived using 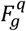 as a query vector and 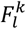 as an index vector, within the context of the adaptive multi-granularity fusion module, as illustrated in Fig. 8. The resulting correlation was calculated as follows:

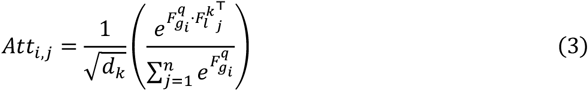

where *Att*_*i,j*_ represents the attention weight quantifying the alignment between the *i*-th voxel feature in the global query vector 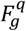 and the *j*-th voxel feature in the local index vector 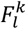. Correlation scores between different local features were then computed using a scaled dot-product operation. To stabilize the attention distribution and prevent over-concentration in specific local regions, a scaling factor of 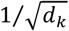 was applied, where *d*_*k*_ denotes the dimension of an individual attention head, corresponding to the number of channels within 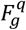. After computing the similarity matrix *Att*_*i,j*_, it is mapped back into the original feature space, encompassing both the global content vector space 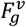 and the local content vector space 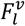. Specifically, the given similarity matrix *Att* undergoes direct propagation into a Softmax function, to produce a sparse feature distribution for the local fine-grained branch 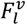. This distribution is then subjected to matrix multiplication with 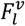, followed by convolution operations to further refine the feature distribution. This sequence can be represented by:

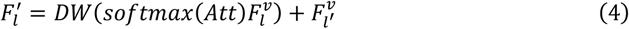

In the case of a global coarse-grained branch 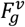, the matrix *Att* requires transposition to determine the similarity score between individual feature points and the query feature. Subsequently, matrix multiplication with 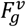 is performed and the resultant product is processed using deep separable convolutions to enhance the smoothness of feature distributions. In addition, a residual technique is employed to preserve the integrity of information and mitigate issues with vanishing gradients. Ultimately, this module yields the following output:

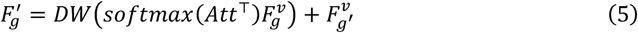

#### Tracing Boundary Contrast Learning

Most existing 3D segmentation algorithms (*22–24*) neglect the explicit segmentation of structural boundaries. However, some algorithms strive to improve boundary delineation using attention mechanisms (*25,26*). Despite these efforts, there remains a lack of comprehensive and targeted research focused on evaluating the influence of boundary regions on segmentation outcomes. To address this issue, we propose meticulously aligning boundary information from manual annotations with model segmentation outputs, thereby introducing an innovative boundary contrast learning framework.

Previous contrastive learning methodologies have predominantly been applied at the image level. For example, positional techniques (*27*) extend 3D images to 2D sequences to determine spatial relationships between positive and negative samples as part of contrastive learning. The Mask Transfiner model (*28*) adopts contrastive learning within image patches, rather than at the image level, utilizing pseudo-labels to generate patches with distinct semantic categories and forming positive and negative sample pairs. However, these strategies are inadequate for high-resolution segmentation of the amygdala, which exhibits proximity with same-type pixels throughout a 3D image. It requires the separation of pixel types and the clustering of similar pixels based on individual labels. Here, boundary pixels were used to construct positive and negative sample pairs to maximize distinctions between boundary pixels and adjacent non-relevant pixels, as illustrated in Fig. 8. Specifically, the local neighborhood around the boundary was designated as a subset, establishing a contrast learning strategy for overlapping 3D patch areas distributed throughout 3D space. Kernels adjacent to the boundary were then used to select the dimensions of 3D patch spaces in each subset. Manually segmented amygdala data thus served as a basis for constructing both positive and negative samples. The InfoNCE information maximization algorithm (*29*) was then used to calculate the disparity between positive and negative samples in the latitude space on a hypersphere. The kernel size was set to 5 × 5 × 5 (based on the anatomical dimensions of the amygdala) and each subset formed positive and negative samples within this specified area. As illustrated in Fig. 8, the model was specifically designed to focus on and exclusively compute boundary voxels. Initially, all boundary regions *B*_*l*_ were considered in manual annotations. For each voxel *x*_*i*_ in *B*_*l*_, the sampling scope for positive and negative voxels was then confined to the local neighborhood *N*_*i*_. Under this constraint, positive samples *x*_*j*_ ∈ *N*_*i*_ ∧ *l*_*i*_ = *l*_*j*_ were identified for boundary voxels *x*_*i*_, while any remaining neighborhood voxels were treated as negative samples *x*_*j*_ ∈ *N*_*i*_ ∧ *l*_*i*_ ≠ *l*_*j*_. In this paper, *j* denotes sample voxels within the neighborhood *N*_*i*_ corresponding to the voxel *i*, with *l*_*i*_ and *l*_*j*_ representing the voxel categories for *x*_*i*_ and *x*_*j*_, respectively. Consequently, the boundary voxel contrast learning algorithm enhanced feature discernment for structurally organized boundary voxels, thereby establishing a target for contrast optimization within boundary voxels, which is crucial for enhancing the segmentation of boundary regions. Specifically, for the voxels *x*_*i*_ within boundary regions *B*_*l*_, learned representations were incentivized to be more congruent with voxel features from the same category, while being less congruent with features of disparate categories. This process can be described as follows:

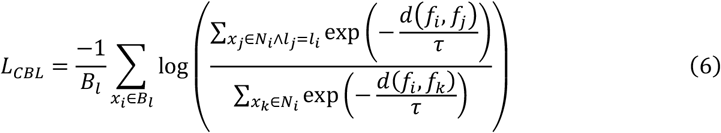

where *f*_*i*_ represents the characteristics of *x*_*i*_, d(·,·) denotes the distance function, and *τ* is a temperature coefficient used in contrastive learning. The numerator captures the similarity between positive instances, while the denominator reflects the similarity between positive and negative instances. In a contrastive loss framework, the objective is to maximize similarity among positive pairs while minimizing similarity between positive and negative pairs. Consequently, increasing intra-class similarity (bringing positive instances closer) while decreasing inter-class similarity (pushing positive and negative instances further apart) leads to a lower loss. The temperature coefficient *τ* is a predefined hyperparameter that controls model’s sensitivity to negative samples. Larger values of *τ* flatten the distribution of *d*(*f*_*i*_, *f*_*j*_), causing the model to treat all negative samples more uniformly, which reduces learning efficacy. Conversely, smaller values of *τ* emphasize more difficult negative samples, some of which may be mislabeled positive samples, leading to false negatives. This approach limits model convergence and restricts generalization capabilities. Therefore, selecting an optimal temperature coefficient is crucial to balancing learning efficiency and discrimination ability.

#### SAM Consistency Module

The SAM framework was used to develop a consistency strategy to mitigate discrepancies arising from the scanning of different subjects. This strategy is comprised of three integral components: a volume feature encoder, a 3D prompt encoder, and a volume mask decoder. The principal contributions of each can be specifically delineated as follows. First, an SAM is introduced, which is specifically designed for amygdala segmentation tasks. Unlike existing approaches to improving SAMs, the novelty of our model lies in its comprehensive consideration of factors such as the volume, shape, and relative position of the amygdala. This process eschews conventional slice-by-slice feature extraction in favor of a holistic total volume approach. The model also establishes interslice associations, which are exceptionally well-suited to structures with intricate core regions. In contrast to traditional algorithms, this technique provides a more comprehensive representation of anatomical structures, particularly for smaller entities.

Second, the incorporation of a 3D prompt encoder ensures the preservation of prior spatial position information and 3D spatial features provided by the user. In contrast to 2D prompt encoders, this strategy provides more precise relative position cues in multidimensional space. The effective merging of 3D cue information with embedding features derived from a volume feature encoder is then facilitated by a novel volume mask decoder. This decoder harnesses interaction mechanisms between prior cue and MRI embeddings, facilitating seamless integration. The fusion mechanism then adeptly captures nuanced details within the feature space, offering a robust tool for the precise depiction of anatomical structures. SAMs constitute an interactive segmentation methodology predicated on iterative operations. This framework enhances the representational capacity of the model through the incorporation of positional cues and pre-segmentation data, thereby facilitating continuous optimization of segmentation accuracy via multiple interactive iterations. Notably, the accuracy of existing SAMs is suboptimal when applied directly to medical imaging tasks (especially amygdala segmentation). To address this issue, the structure of the proposed 2D SAM has been meticulously customized for amygdala segmentation. This includes the introduction of an advanced SAM, which is comprised of three integral encoders used for volume features, 3D cues, and volume masks.

The function of the Volume Feature Encoder (VFE) is to transform input T1w structural images into an MRI embedding vector. Unlike the 2D encoder used in traditional SAMs, this process allows simultaneous consideration of volume, shape, and relative position, thereby offering a more comprehensive representation of anatomical structures to enhance the processing of complex regions surrounding the core area. A 3D prompt encoder was also used to encode prior information, incorporating interactive cues and existing amygdala segmentation results. The 3D prompt encoder consists of two components: a click-cue branch and a presegmentation cue branch. The click-cue branch converts user-provided cue points into learnable location embedding features to more precisely delineate amygdala location. The pre-segmentation prompt encoder uses the last click made during interactive segmentation as input. Alternatively, it provides pre-segmentation results to the model, thereby prompting adjustments and optimization of the current segmentation result. As such, a volume mask detector (VMD) was developed to effectively integrate information from the prompt encoder with embeddings produced by the feature encoder. This VMD also facilitated the retrieval and refinement of information pertaining to the amygdala, ensuring a seamless alignment of cue feature information and initial feature encoder output.

The VMD is a pivotal component to exhaustively explore key characteristics within the amygdala, thereby enhancing model efficacy in intricate clinical scenarios. It aims to combine MRI embedding vectors (produced by the volume feature encoder) with userprovided prompt inputs (used to determine the correlation between them), culminating in precise predictive results. The 3D mask decoder generates MRI and cue embeddings using a multi-layered fusion 3D feature encoder, facilitating multi-level and multiperspective information extraction. This comprehensive fusion mechanism enables a more complete assimilation of MR images and improves the differentiation of anatomical structures, thereby increasing predictive precision. In conclusion, the decoder generates precise predictions of specific structures or lesions within images, offering crucial support for clinicians in making accurate diagnoses and devising treatment strategies.

### Training Configuration

#### Implementation of Loss Function

We implemented an advanced deep-supervised strategy to formulate the loss function. Specifically, this involved the integration of supervisory signals in each intermediate and terminal module, thereby facilitating enhanced gradient propagation and optimization of network training efficiency. Two distinct loss functions were incorporated for each supervisory signal, namely, binary crossentropy (BCE) and Dice loss. The overall loss function was constructed as follows:

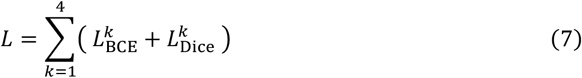

An adapted version of BCE loss was employed to address issues with imbalances between positive (amygdala) and negative (other tissue) samples. Hereafter, the superscript *k* is excluded for simplicity, while L_*extBCE*_ is defined as follows:

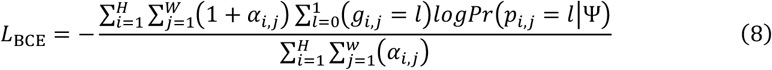

where *l* ∈ (0, 1) denotes two label categories. The variables *p*_*i,j*_ and *g*_*i,j*_ represent predicted and ground truth values, respectively, at the spatial coordinate (*i, j*) within a CT image characterized by a width of *W* and a height of *H*. The symbol Ψ encompasses all model parameters and *Pr*(*p*_*i,j*_ = *l*) signifies the predicted probability. The term *α*_*i,j*_ ∈ [0, 1] describes the pixel-specific weight and can be expressed as:

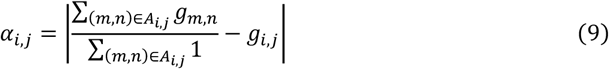

where A*i,j* denotes the region adjacent to pixel (*i, j*) and | · | signifies the calculation of absolute value. Furthermore, Dice loss was defined as:

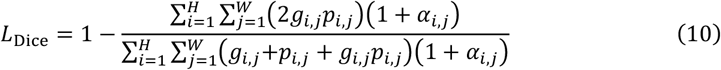

#### Implementation Details

The model was implemented using the PyTorch framework and trained on a full capabilities of the Tesla A10 GPU to optimize computational efficiency. The AdamW optimizer (*30*), a common choice within the transformer architecture domain, was used for routine updates of network parameters. The learning rate was empirically set to 1e^−4^, balancing training stability and convergence speed, with a weight decay of 1e^−4^. Memory consumption was mitigated by designating a batch size of 1, with a training process encompassing 100 iterations. This configuration ensured the model underwent sufficient training cycles to achieve optimal performance and effectively discern feature representations.

### Evaluation Metrics

Model performance was assessed using standard medical image segmentation metrics, including region overlap measures (Dice Similarity Coefficient and Jaccard Similarity), boundary distance measures (Hausdorff Distance (HD95), Average Surface Distance (ASD), and Average Symmetric Surface Distance (ASSD)), volumetric comparison (Relative Volume Difference, RVD), and classification metrics (Precision and Recall) (*31*).

**The Dice similarity coefficient (Dice)** measures the similarity between a predicted binary region (P) and a ground truth (GT). It is formulated as:

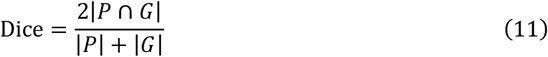

where *P* and *G* denote voxel sets for the segmented area and GT, respectively. The term | · | denotes a cardinality computation operation, which provides the number of elements in a set.

**Jaccard Similarity (JS)** is defined as the area of intersection between predicted segments and labels, divided by the area of union between the two, expressed as:

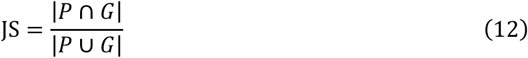

**Hausdorff Distance (HD)** measure the maximum surface distance distribution between the predicted segmentation result and the ground truth. HD95 is an improvement on the Hausdorff distance that reduces the impact of outliers by taking the 95% quantile.

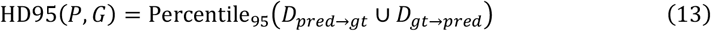

**Average Surface Distance (ASD)** is used to measure the surface distance difference between the predicted segmentation result and the ground truth.

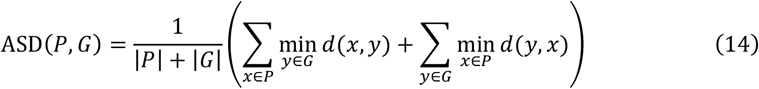

**Average Symmetric Surface Distance (ASSD)** is a commonly used symmetric surface distance metric in medical image segmentation evaluation. It measures the surface distance difference between the predicted segmentation result and the ground truth, while considering the bidirectional distance from prediction to GT and from GT to prediction.

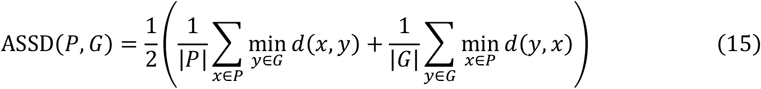

**Precision** is a commonly used evaluation metric that measures the proportion of positive predictions in the predicted segmentation results that are correct, especially for unbalanced datasets.

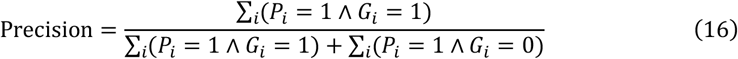

**Recall** is another important evaluation metric that measures how many positive pixels in the true segmentation are correctly predicted as positive. It reflects the coverage of the model for positive samples.

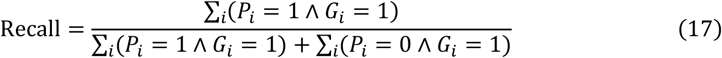

**Relative Volume Difference (RVD)** is one of the commonly used evaluation indicators in medical image segmentation tasks, which is used to measure the volume difference between the predicted segmentation result and the GT.

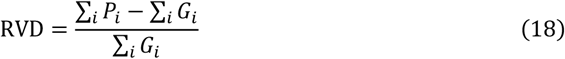

## Supplementary Materials

### This PDF file includes

Table S1, S2

## Funding

This work was supported by The STI 2030 - the major projects of the Brain Science and Brain-Inspired Intelligence Technology 2021ZD0200500 (X-N.Z.)

Start-up Funds for Leading Talents at Beijing Normal University (X-N.Z.)

National Natural Science Foundation of China 82102134, 82322035, 62273076 (X-N.Z.)

Key-Area Research and Development Program of Guangdong Province 2019B030335001(X-N.Z.)

Beijing Municipal Science and Technology Commission Z161100002616023, Z181100001518003 (X-N.Z.)

Major Project of National Social Science Foundation of China 20&ZD296 (X-N.Z.)

CAS-NWO Programme 153111KYSB20160020 (X-N.Z.)

National Basic Research (973) Program 2015CB351702 (X-N.Z.)

Chinese Academy of Sciences Key Research Program KSZD-EW-TZ-002 (X-N.Z.)

International Collaboration of National Natural Science Foundation of China 81220108014 (X-N.Z.)

China Postdoctoral Science Foundation 2023M740301 (Q.Z).

## Author contributions

Conceptualization: H-J.H., X-N.Z.

Methodology: B.D., Q.Z., P.G.

Investigation: Q.Z., B.D., P.G.

Resources: X-N.Z., H-J.H.

Data curation: Q.Z., P.G.

Visualization: B.D., P.G.

Supervision: H-J.H., X-N.Z.

Writing—original draft: B.D., Q.Z.

Writing—review and editing: All authors contributed to the critical review and editing of the manuscript.

## Data and materials availability

All data needed to evaluate the conclusions in the paper are present in the paper and/or the Supplementary Materials. The manually segmented amygdala datasets from the Chongqing cohort of the Chinese Color Nest Cohort (CKG-CCNC), used for model training and internal validation, are publicly available via the National Science Data Bank (https://doi.org/10.11922/sciencedb.01299). The original MRI scans and corresponding demographic information (age and gender) are accessible from the Chinese Color Nest Project - Lifespan Brain-Mind Development Data Community (https://ccnp.scidb.cn/en). External validation was conducted using the Zhejiang University Multi-center Traveler Cohort (ZJU-MTC), which is available on Figshare (https://figshare.com/articles/dataset/Multicenter_dataset_of_multishell_diffusion_magnetic_resonance_imaging_in_healthy_traveling_adults_with_identical_setting/8851955/6). Due to data-sharing agreements and institutional review board restrictions, the BNU-CAC dataset is not publicly available. All source code used for training, evaluation, and visualization is publicly available at the Science Data Bank (https://ccnp.scidb.cn/en). For detailed usage instructions, please contact the first author.

## For the Chinese Color Nest Consortium (CCNC)

Xi-Nian Zuo^1,2,3,4,7,18^, Ning Yang^1,2,3,4^, Zhe Zhang^5^, Ye He^6^, Hao-Ming Dong^1,4,8^, Lei Zhang^9^, Xing-Ting Zhu^2^, Xiao-Hui Hou^7^, YinShan Wang^1,3^, Quan Zhou^1,2,3^, Zhu-Qing Gong^1,2,3^, Li-Zhi Cao^2,4^, Ping Wang^1^, Yi-Wen Zhang^2^, Dan-Yang Sui^20^, Ting Xu^21^, GaoXia Wei^2,4^, Zhi Yang^2,4^, Lili Jiang^2,4^, Hui-Jie Li^2,4^, Ting-Yong Feng^11^, Antao Chen^11^, Jiang Qiu^11^, Xu Chen^11^, Xun Liu^2,4^, Ke Zhao^2,4^, Yi Du^2,4^, Min Bao^2,4^, Yuan Zhou^2,4^, Yan Zhuo^14^, Zhentao Zuo^14^, Li Ke^1^, Fei Wang^22^, Zhi-Xiong Yan^7^, Xue-Quan Su^7^, F.Xavier Castellanos^23^, Michael Peter Milham^21,24^, Yu-Feng Zang^25^

^1^State Key Laboratory of Cognitive Neuroscience and Learning, Beijing Normal University, Beijing, 100875, China.

^2^Department of Psychology, University of Chinese Academy of Sciences, Beijing, 100049, China.

^3^Developmental Population Neuroscience Research Center, International Data Group/McGovern Institute for Brain Research, Beijing Normal University, Beijing, 100875, China.

^4^Key Laboratory of Behavioural Science, Institute of Psychology, Chinese Academy of Sciences, Beijing, 100101, China.

^5^College of Education, Hebei Normal University, Shijiazhuang, 050024, China.

^6^School of Artificial Intelligence, Beijing University of Posts and Telecommunications, Beijing, 100876, China.

^7^Laboratory of Cognitive Neuroscience and Education, School of Education Science, Nanning Normal University, Nanning, 530299, China.

^8^Changping Laboratory, Beijing, 102206, China.

^9^School of Government, Shanghai University of Political Science and Law, Shanghai, 201701, China.

^10^School of Psychology, Research Center for Exercise and Brain Science, Shanghai University of Sport, Shanghai, 200438, China.

^11^Faculty of Psychology, Southwest University, Chongqing, 400715, China.

^12^NYU-ECNU Institute of Brain and Cognitive Science at New York University Shanghai, Shanghai, 200062, China.

^13^Faculty of Arts and Science, New York University Shanghai, Shanghai, 200122, China.

^14^State Key Laboratory of Brain and Cognitive Science, Institute of Psychology, Chinese Academy of Sciences, Beijing, 100101, China.

^15^School of Psychology and Cognitive Science, East China Normal University, Shanghai, 200062, China.

^16^Department of Psychology, Renmin University of China, Beijing, 100872, China.

^17^Beijing Key Laboratory of Learning and Cognition, School of Psychology, Capital Normal University, Beijing, 100048, China.

^18^School of Education, Hunan University of Science and Technology, Hunan Xiangtan, 411201, China.

^19^National Basic Science Data Center, Beijing, 100190, China.

^20^Flight Research Department, Aviation University of Air Force, Jilin Changchun 130012, China.

^21^Center for the Developing Brain, Child Mind Institute, New York, NY, USA.

^22^Affiliated Nanjing Brain Hospital, Nanjing Medical University, Nanjing 210000, China.

^23^Department of Child and Adolescent Psychiatry, New York University Grossman School of Medicine, New York, NY 10016, USA.

^24^Nathan S. Kline institute for Psychiatric Research, Orangeburg, New York, NY 10962, USA.

^25^Institute of Psychological Sciences, Hangzhou Normal University, Hangzhou 311121, China.

